# Optimal Control and the Dynamics of Ancestral Lineages

**DOI:** 10.64898/2025.12.04.692409

**Authors:** Colin LaMont

## Abstract

In resource-constrained populations, the growth rate of an allele is coupled to the frequency of its competitors. Since these frequencies are unobserved, exact inference of evolutionary parameters demands integrating over the full ensemble of possible frequency histories. This high-dimensional path integral is computationally prohibitive, and standard phylogenetic approaches avoid its evaluation by treating lineages as unconstrained birth-death processes. However, ignoring the carrying capacity comes at a cost: it necessitates an abundance of free parameters to describe simple allelic replicator dynamics and implicitly assumes an incorrect drift model.

We propose a new approach by defining the *fitness potential* (Ψ)— a dynamical quantity formally equivalent to the logarithm of reproductive value. By exploiting a duality between evolutionary dynamics and stochastic optimal control, we show that Ψ allows for the analytic marginalization of the unknown population history. This transformation yields a tractable likelihood function that strictly enforces competitive constraints while accounting for the information loss due to genetic drift in finite populations. We demonstrate that this framework enables parsimonious, site-specific inference of selection coefficients from phylogenetic trees. This dual-control framework shows ancestral lineages behave as optimal agents: they do not simply climb local fitness gradients, but “anticipate” future selection by navigating a global landscape of reproductive value.

## 1 Introduction

The fitness landscape is a foundational concept in evolutionary genetics, mapping genotypes to their reproductive success. However, in any finite environment, success is relative: an allele’s growth is constrained by competition with the rest of the population. Mathematically, this means the birth and death rates of a specific genotype *g* (denoted *β*_*g*_ and δ_*g*_) are linked not to absolute fitness *f*_*g*_, but to the fitness relative to the population mean, 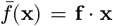. Consequently, calculating the likelihood for fitness parameters requires continuous knowledge of this mean fitness term, which depends on the frequency of every allele in the population state **x**. Capturing the rich physics of evolution (for example, clonal interference and competitive exclusion) therefore requires integrating over the full ensemble of possible population histories, i.e. evaluating a path integral over the trajectory of allele frequencies **x**(*t*). This task faces a severe curse of dimensionality. As the number of genetic variants or time points increases, the space of possible population trajectories grows exponentially, making the exact path integral computationally prohibitive.

To achieve tractability, standard phylogenetic approaches (like birthdeath skyline models [6, 47]) avoid this high-dimensional integration by decoupling the lineages. They treat each lineage’s growth rate as an independent, piecewise-constant parameter to be inferred, effectively ignoring the zero-sum constraint that couples lineage growth to the population state **x**(*t*). While this avoids the statistical circularity of conditioning on an inferred trajectory (a “double use of data”), it comes at the cost of model parsimony: it requires an explosion of free parameters to describe dynamics that are physically generated by a small number of selection coefficients acting within the governing constraint of the carrying capacity. We propose a framework for the selective pressure on lineages that strictly respects a finite carrying capacity, enabling parsimonious inference in these biologically relevant regimes. To achieve this, we exploit a formal duality with stochastic optimal control theory, drawing inspiration from methods where log-ratios are used to measure the “value” of a state. We propose a similar quantity for population genetics to measure the reproductive value of an allele [17, 19, 20, 3]. This quantity, the fitness potential Ψ_*g*_(**x**), is defined as the log-ratio of an allele *g*’s frequency in the ancestral pool, *w*_*g*_(**x**), relative to its frequency in the contemporaneous living population, *x*_*g*_:

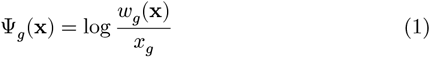

Intuitively, this potential measures the enrichment of an allele among ancestors relative to the population in which they lived.

We derive the backward-in-time dynamics of this quantity from standard population-genetic principles. A key result of our analysis is that, in the weak-selection limit, this potential approximately decomposes into a frequency-independent component specific to each allele (*ψ*_*g*_) and an allele-independent component shared by all possible lineages (*φ*(*x*)):

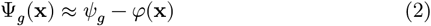

This decomposition is the mechanism that allows tractable inference. By isolating the complex, frequency-dependent population dynamics within a shared term (*ϕ*(**x**)), we can effectively marginalize out these unobservable histories. The relevant information for comparative success is retained in the much simpler, allele-specific term (*ψ*_*g*_), which is governed by an ordinary differential equation (ODE) rather than a curse-of-dimensionality-afflicted partial differential equation (PDE).

We explore the computational and conceptual implications of this simplification. First, we show how this formalism provides a new lens on adaptation in dynamic environments, revealing a formal duality with optimal control theory. Second, we benchmark the approximation against exact solutions in biallelic systems to establish its valid parameter regimes. Finally, we leverage this validated theory to develop an efficient likelihood framework for inferring time-varying selection directly from phylogenetic trees.

## 2 Methods

### 2.1 Model setup and definitions

We model the evolution of a population using the language of a standard Wright-Fisher diffusion or Moran process. Let **x** = (*x*_1_, *x*_2_, …, *x*_*k*_) denote the vector of genotypic frequencies, where *x*_*g*_ represents the relative fraction of genotype *g* in the living population such that Σ*x*_*g*_ = 1(Figure 1, Panel A). This vector encodes the population state, **x**(*t*) and evolves forward in time according to a stochastic process driven by mutation (rate matrix *μ*), selection (fitness vector **f**), and random genetic drift (scaled by population size *N*_*e*_). As we consider haploid populations (e.g., viral strains), we use the terms *genotype* and *allele* interchangeably to refer to the specific genetic state of a lineage.

**Figure 1.**
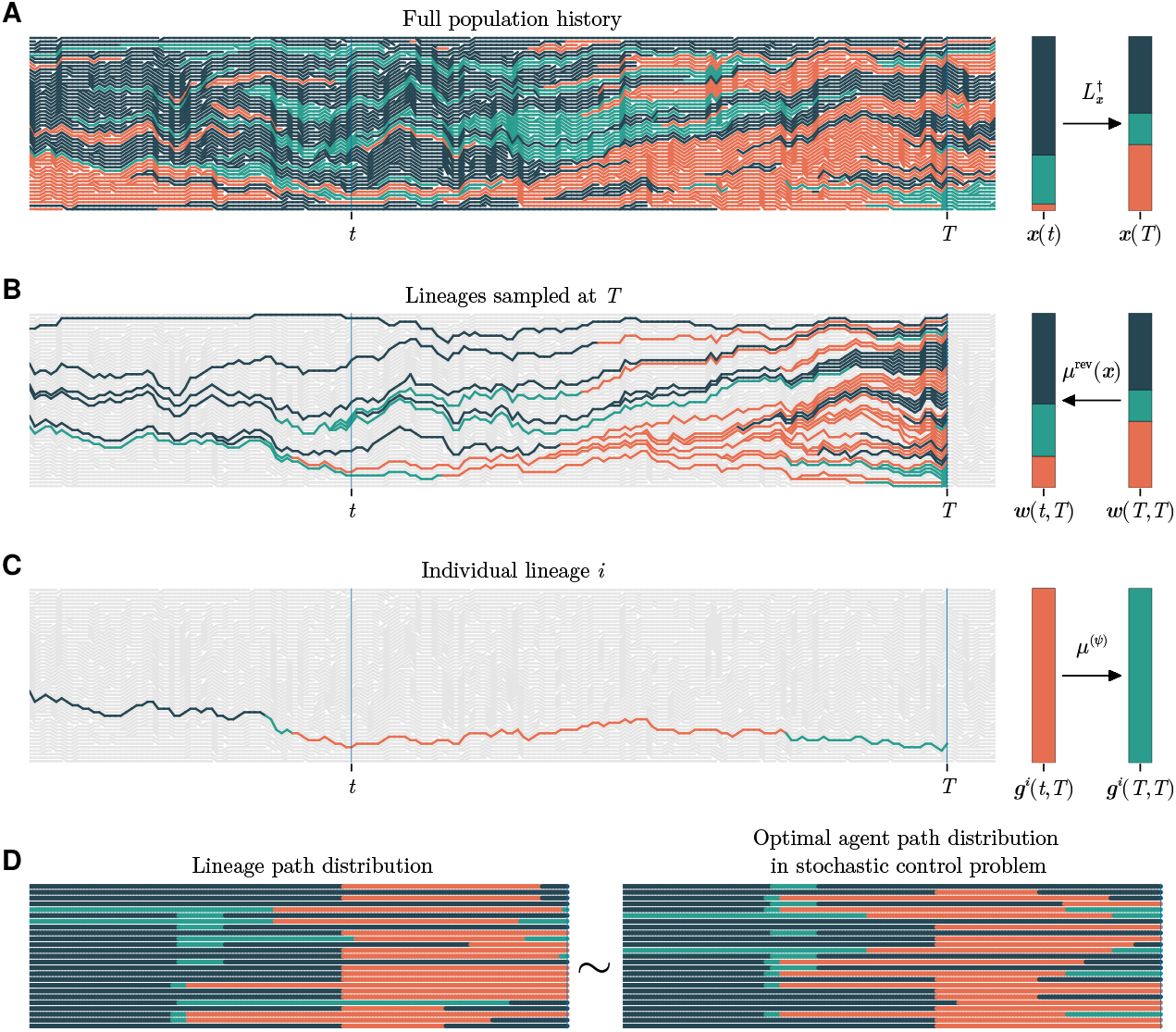
The correspondence between ancestral lineage dynamics and optimal control. (A) The full population history evolves forward in time from *t* to *T*, driven by mutation, selection, and drift (mathematically described by the forward Fisher-Kimura generator 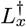), resulting in the observable population state *x*(*t*) (B) Conditioning on survival to the sampling time *T* reveals the ancestral lineages (colored paths). The ancestral frequency distribution *w*(*t, T*) represents the probability that an ancestor at time *t* has a specific genotype, evolving backward according to the *x*-conditioned operator *μ*^rev^(**x**). (C) A single surviving lineage *i* traces a discrete path *g*^*i*^(*t, T*) through genotypic space. While individuals in the full population generate mutations according to *μ*, the surviving lineage appears to follow a biased mutational operator *μ*^(*ψ*)^, where the bias is driven by the fitness potential *ψ*. (D) The distribution of these biological lineage paths (left) is statistically identical to the distribution of optimal agent trajectories in a dual stochastic control problem (right). This duality implies that surviving lineages appear to have “anticipated” future selective pressures by following an optimal control policy.

#### The Ancestral Lineage

Our primary goal is to understand the dynamics of a single lineage traced backward from a sampled individual. Unlike the general population, which is shaped by mutation, selection and genetic drift, ancestral lineages have been shaped by an additional factor: the retrospective condition of having successfully left descendants. Consider an experiment where we sample a single individual at the present time *T*. We wish to trace its ancestry backward to time *t* < *T*. Let *w*_*g*_(**x**, *t, T*) be the probability that the ancestor at time *t* had genotype *g*, given the population state was **x** at *t*. As depicted in Figure 1, Panel B, at *t* = *T*, the ancestor will be identical to the sampled individual, implying the boundary conditions that **w**(**x**, *T, T*) = **x**. We derive the dynamics of **w**(**x**, *t, T*) as *t* recedes into the past. We will then perform a change of variables to find the dynamics of the 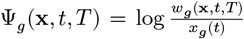, our key quantity of interest.

We consider a fixed sampling time *T*, and suppress the *T* dependence in our notation. Furthermore, where the time-dependence *t* is clear from context, we will write *w*_*g*_(**x**), and Ψ_*g*_(**x**) for brevity.

### 2.2 Backward evolution of the ancestral frequency

We begin with the dynamics of the ancestral frequencies **w**(**x**, *t*), defined as the probability that an ancestor at time *t* has genotype *g*, given the population state was **x**. Lineage transport between genotypic states is governed by the frequency-conditioned reversed mutational operator, *μ*^rev^. We derive this by applying Bayes’ rule to the mutation flux. In the forward direction, the total flux of individuals mutating from *g*′ to *g* is proportional to the mutation rate 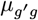 and the frequency of the source *x*_*g*_′. The probability that a specific lineage in state *g* originated in *g*′ is this flux normalized by the abundance of the destination state *x*_*g*_:

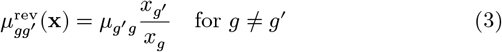

To conserve probability, the diagonal elements are set to the negative sum of the off-diagonals. As we move backward in time, this operator pulls lineages toward historically abundant genotypes. Notably, birth and death events do not appear in this transport term; tracing a surviving lineage backward, coalescent events concentrate the distribution into fewer distinct ancestors, but do not alter the genotype of the lineage itself.

Combining this reversed mutational process with the adjoint Kimura diffusion operator (*L*_**x**_) [27], which moves the **x**-conditioning backward in time, we arrive at the full evolution equation:

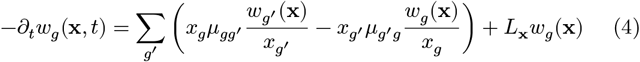

We now perform a change of variables from *w*_*g*_(**x**) to our core object, the fitness potential Ψ_*g*_(**x**) = log(*w*_*g*_(**x**)/*x*_*g*_). Applying this transformation to the backward dynamics in Equation 4 yields the backward-time evolution for the potential

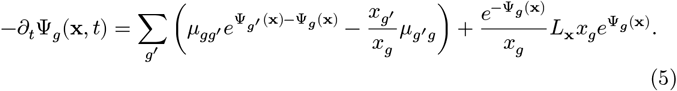

Expanding this 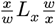 sandwich on the selection, mutation, and drift-diffusion operators (see Appendix B) yields the exact backward evolution for the fitness potential,

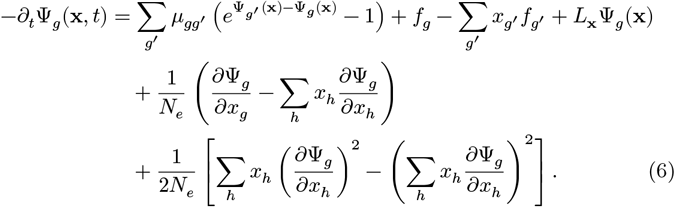

#### 2.2.1 The forward generator for phylogenetics

This backward-time equation for Ψ has a key role in defining the apparent forward mutation rates along a surviving lineage. The apparent forward mutation rates are a central observable quantity in phylogenetics, propagating the likelihood back up the tree in Felsenstein’s pruning algorithm [14]. These rates obey the **w**-conditioned reverse of the **x**-conditioned reverse of the bare mutation operator. From Equation 3, a second application of Bayes rule yields off-diagonal mutation rates

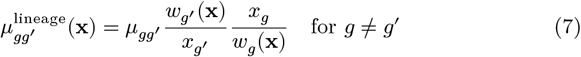

This apparent forward mutation operator only depends on 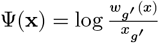. We can thus write the forward mutational operator as a function of Ψ(**x**):

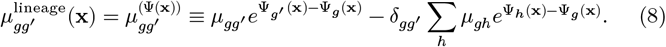

This result provides the bridge between our backward-time-propagating potential and forward-time phylogenetic inference.

### 2.2.2 Approximate splitting between state and frequency dependence

Our boundary condition is that Ψ_*g*_(**x**, *T*) = 0. Ψ_*g*_(**x**) is therefore a (trivially) **x**-independent function at *T*. Frequency dependence is introduced at first order in time through the *g*-independent competition term, Σ_*h*_ *f*_*h*_*x*_*h*_ which enforces normalization, Σ_*g*_ *x*_*g*_ = 1.

We take the ansatz that the frequency dependence, which was trivially allele-independent (zero) at *T*, continues to be allele-independent as we move backward in time. Specifically, we assume we can write Ψ_*g*_(*x*) = *ψ*_*g*_ − *φ*(*x*). The conservation of the number of lineages then implies Σ_*g*_ *w*_*g*_ = 1 so we can express the shared frequency function *φ*(*x*) as a function of the *ψ*_*g*_

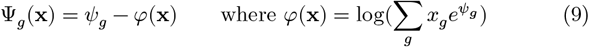

Plugging into Equation 6

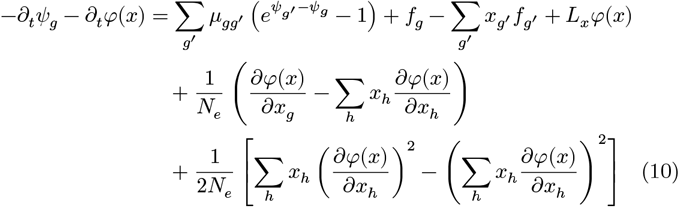

To study the *ψ* evolution we only need consider the *g*-dependent terms on each side of the equation. Importantly, we have a gauge freedom in the definition of *ψ*_*g*_: an allele-independent shift *ψ*_*g*_(*x*) → *ψ*_*g*_ + *φ*′(*x*), shifts *φ*(*x*) → *φ*(*x*) + *φ*′(*x*), and leaves Ψ_*g*_ and all observable dynamics unchanged. However, the choice of *φ*′(*x*) may introduce or remove apparent frequency dependence in *ψ*_*g*_. Choosing a gauge where the frequency coupling is eliminated as much as possible (to first order),

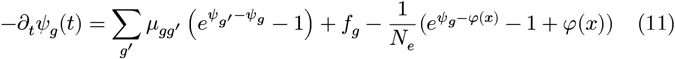

Thus, where the differences between *ψ*_*g*_ and *φ*(*x*) stays relatively small

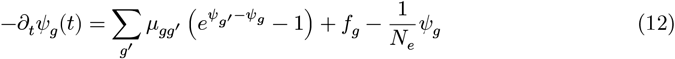

where this last equation holds up to order 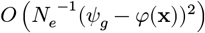. Equation 12 is thus valid in the weak selection limit (eg,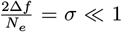) or for selection acting at short timescales 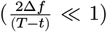, where the magnitude of the fitness potential differences are small.

Knowing *ψ* is enough to construct the forward mutation operator,

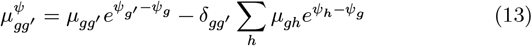

Thus, we have shown that the weak-selection limit, Equation 12 and Equation 13 define the lineage dynamics without reference to the population state **x**.

### 2.3 Connection to optimal control

Our derivation of the frequency-independent fitness potential uncovers a deep duality: where the weak selection limit is valid, the equations governing ancestral lineages are formally identical to those of optimal control theory. (See Appendix A for details about the control theoretic perspective on Equation 12 and Equation 13) In this duality, biological alleles map to agent states, and our fitness potential maps to the control theorist’s “value function”—a measure of maximum future reward. This is not accidental, we defined **w** and Ψ, guided by this duality, specifically to recover and exploit Equation 12, which maps to the Hamilton-Jacobi-Bellman (HJB) equation, the fundamental equation of control theory.

Beyond furnishing us with a tractable ODE and generator for the lineage dynamics, the duality between control and Darwinian evolution provides us a powerful new reference frame for biological intuition. It is common to personify pathogens as clever adversaries—cancer deleting TP53, or bacteria upregulating *β*-lactamase. Reading Equation 13 as a mutational control policy, this evolutionary cleverness can be understood directly as optimal play. Because we only observe lineages that succeeded in surviving to the present, they appear to have followed an optimal policy—anticipating and steering around the challenges posed by mitigation efforts. This anticipation does not arise from biological foresight–it is a mathematical inevitability of conditioning on survival. At the same time, Equation 12 indicates that the capacity for anticipation is limited and exponentially suppressed beyond the population’s coalescence horizon, *N*_*e*_.

## 3 Results

### 3.1 Ancestral stationary distribution of a biallelic system

Let us compare our theory to the exact steady-state biallelic distribution to determine its range of validity in a tractable case. For simplicity, *μ*_12_ = *μ*_21_ and *f*_1_ = 0, *f*_2_ = *f*.

#### The weak-selection approximation compared to selective sweeps

The stationary condition for the fitness potential is given by setting the derivative of Equation 12 equal to zero yields an implicit equation for the potential difference Δ*ψ* = *ψ*_2_ − *ψ*_1_:

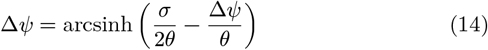

where *σ* = 2*N*_*e*_*f* and *θ* = 2*N*_*e*_*μ*. The behavior of Δ*ψ* has three distinct regimes depending on the diversity *θ* and the selection ratio *f*/*μ*:

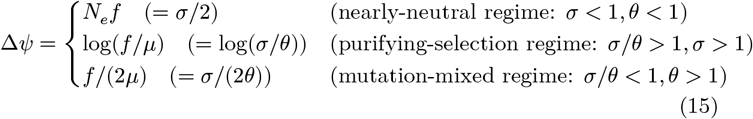

These regimes are sketched in Figure 2.

**Figure 2.**
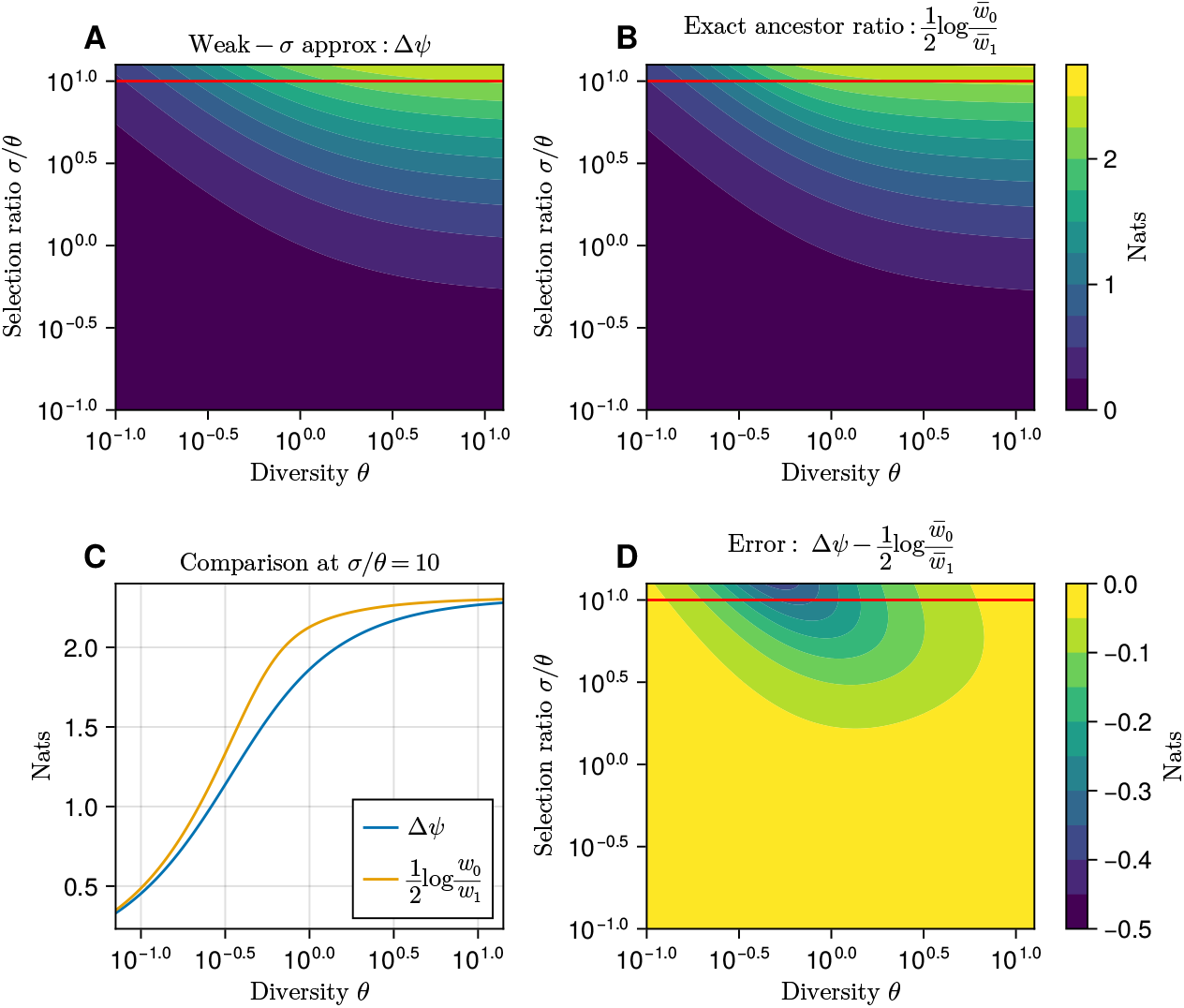
A comparison between the fitness potential estimated from weak selection approximation (Δ*ψ* shown in (A)) and the exact ancestor ratio (B), 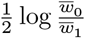, where 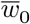 and 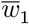 have had their frequency dependence averaged over the equilibrium distribution. (C) A direct comparison between the two measures as a function of the diversity at a fixed selection ratio of *σ*/*θ* = 10. (D) The comparison in the context of the overall error (red line shows the slice in panel C) over the same parameter values as A and B, with the greatest error for intermediate values of *θ* and large selection, corresponding to the regime of strongly-correlated clonal interference.

##### 3.1.1 Comparison to exact results

We now compare this approximation to exact results. We can consider how the weak selection theory compares to stationary ensemble averages,

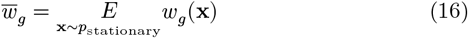

In the biallelic case, the stationary distribution for the population is given by the Wright equilibrium equation:

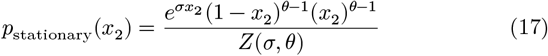

#### Rare Mutation Limit (*θ* ≪ 1)

In this limit, the population toggles between two completely homogenous states. Transitions are governed by the Kimura fixation probability, leading to forward and backward transition rates of 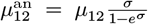 and 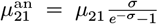. Remarkably, our weak-selection result recovers the correct mutation ratios and stationary distribution even for arbitrarily strong selection in this limit:

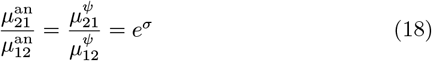

This robustness is surprising, as the linearizing assumptions used to derive our approximation would be expected to fail under strong selection. The mathematical mechanism for this continued validity outside the weak-selection regime remains an open question, suggesting the frequency averaging can potentially recover linearity that is lost when strictly **x**-conditioning population ensembles.

#### Clonal interference regime (Intermediate *θ*)

To study regimes where the weak-selection result may break down, we must calculate the exact stationary ancestral distribution. While complex analytical results exist [13], a simpler approach is to solve our equation for *w*_*g*_(**x**) (Equation 4) numerically using the Galerkin method with the necessary boundary conditions *w*_1_(*x*_1_ = 1) = 1 and *w*_2_(*x*_2_ = 1) = 1 (i.e. the ancestor must be the type of the fixed allele). Comparing these exact numerical results to our approximation (Figure 2) confirms that our theory is accurate at the extreme limits of all three regimes. The greatest errors occur, as expected, at intermediate values of *θ* combined with strong selection, where clonal interference is most significant.

### 3.2 Anticipatory lineage dynamics

We now apply our formalism to dynamic, non-equilibrium scenarios, focusing on the biologically relevant case of time-varying fitness with constant mutation rates. We numerically solve the fitness potential ODE and use it to compute the mutation propagator. We then compare these theoretical predictions to mutation rates observed along lineages in stochastic simulations. We deliberately choose parameter regimes where our weak-selection approximation begins to break down, demonstrating that even in these challenging cases, theory captures the essential qualitative features of the ancestral process.

These examples help illustrate a counter-intuitive phenomenon: ancestral lineages appear to anticipate future selective pressures. As we have emphasized, this is not a violation of causality, but a natural mathematical consequence of conditioning a lineage’s history on its future survival up to the sampling time *T*.

#### 3.2.1 Fitness pulse

We consider a brief selection event between 1.6 and 1.8 *N*_*e*_, which is negative for allele 1 and positive for allele 2. We might imagine this as a brief intervention, such as antibiotic therapy acting on existing gut microflora or an anticancer agent briefly applied to a tumor. The mutational process is symmetric *μ*_12_ = *μ*_21_ = *μ* = 0.1/*N*_*e*_. The fitness function is

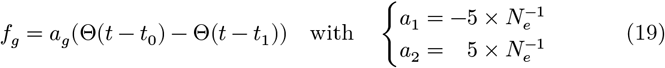

The intensity of the fitness pulse is given in units of *N*_*e*_ and if applied continuously would be associated with an enormous selection coefficient *σ* = 20. Tracing a lineage backward in time sampled at time 2*N*_*e*_, after the pulse is over, it experiences a selection potential given by the blue lines in Figure 3. We see that the mutational operator acting along the lineage, *μ*^*ψ*^ governed by *ψ*, becomes increasingly biased prior to the selection pulse.

**Figure 3.**
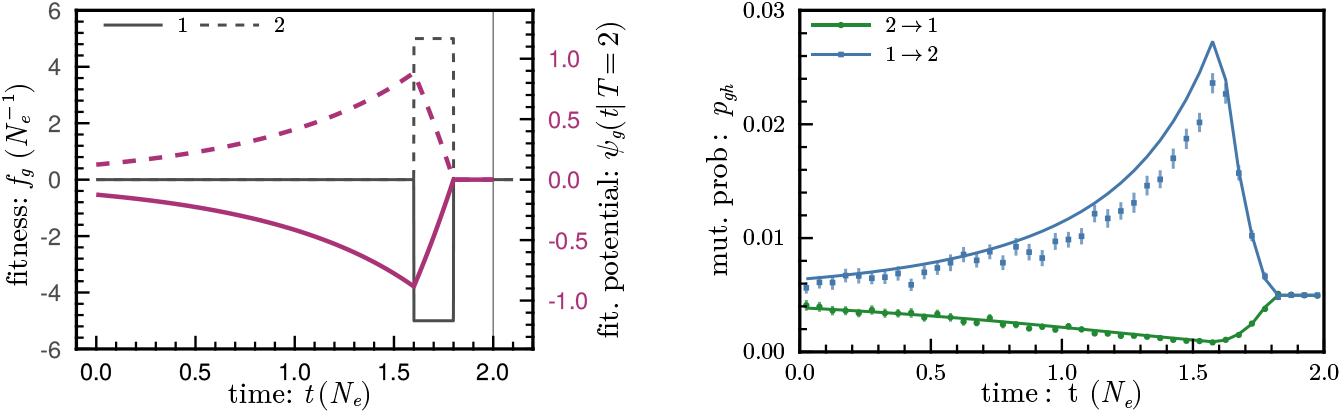
Brief strong selection. We consider a brief selection event between 1.6 and 1.8 *N*_*e*_ on a biallelic population with *θ* = 0.1, balanced mutation rates. Allele 1 experiences negative selection (solid lines) and allele 2 experiences positive selection (dashed lines). Tracing a lineage sampled at time 2*N*_*e*_ backward in time, it experiences a selection potential given by the purple lines. As a result the mutational operator acting along the lineage, *μ*^*ψ*^ governed by *ψ*, is biased before the selection pulse. The mutational process along a lineage is anticipatory in that it occurs before the selection event. (right) The mutation probability, observable in the number of transitions along the lineage over discrete intervals of *N*_*e*_/20 recapitulates the predictions of theory. Since *θ* = 0.1, this predicts a neutral mutation rate of *p*_*gg*_′ = 5 × 10^−3^ observed after the selection pulse is over. During and prior to the selection pulse, the fitness potential becomes much higher for allele 2 than for allele 1 and mutations into the positively selected allele are enhanced, and the negative allele are reduced.

The mutation rates along a lineage appear to be subject to an optimal controller trying to drive the lineage into the positively selected state 2. This control slowly turns on prior to the fitness pulse, and turns back off as we approach the end of the pulse. In this sense, the process as anticipatory, though this anticipation is purely the result of *post-hoc* conditioning a lineage on having surviving offspring at the sampling time *T* (in Figure 3 *T* = 2*N*_*e*_). Looking at the distribution of mutations along a lineage as shown in Figure 3, we confirm that the mutational biases decay as the fitness potential decays exponentially at rate 1/*N*_*e*_, showing that the coalescent time-scale *N*_*e*_ sets the discounting factor in the dual control problem, as we concluded from studying the steady state limit of the biallelic system.

#### 3.2.2 Seasonal adaptation in viral infections

Suppose we have two phenotypes of an influenza strain, one of which is summer-adapted, and one of which is winter-adapted. We can write the fitness function as

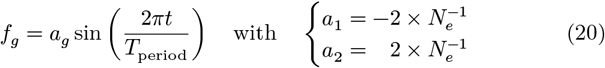

In Figure 4, *T*_period_ = 2.3*N*_*e*_. Considering the living population, obeying the diffusion equation with a time varying flux, observed phenotypes in the living population are lagging the fitness effects since the living phenotype distribution is forward integral of the Kimura equation. Thus the summeradapted strains are, broadly speaking, most dominant in the fall after the cumulative effect of the entire summer of fitness pressure.

**Figure 4.**
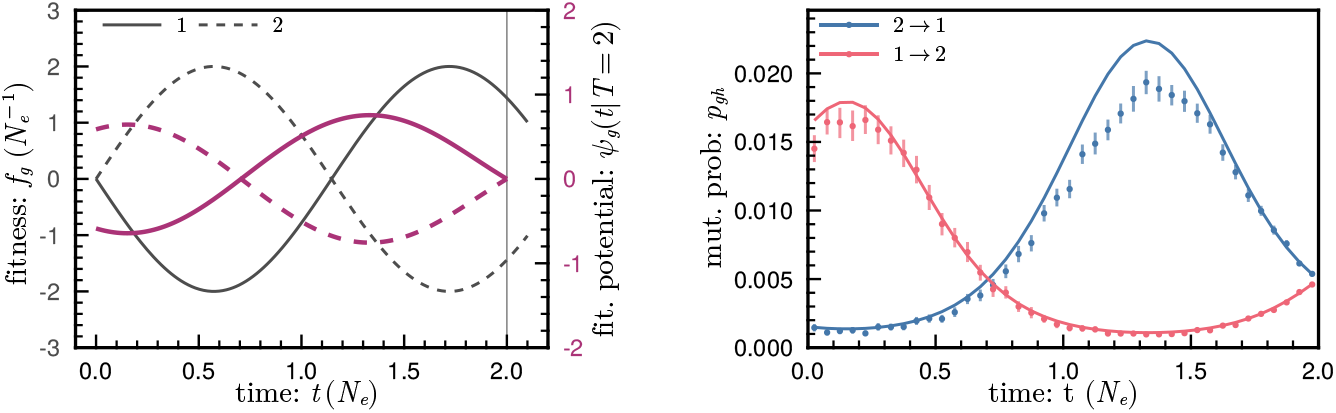
Sinusoidally varying selection. We simulate the effect of a sinusoidal selection (gray lines) event between 1.6 and 1.8 *N*_*e*_ on a biallelic population with *θ* = 0.1, and balanced mutation rates. Tracing a lineage sampled at *T* = 2.0 backward in time, it experiences a selection potential given by the purple lines. As a result the mutational operator acting along the lineage, *μ*^*ψ*^ governed by *ψ*, is biased before the selection pulse. The mutational process is anticipatory in that it occurs before the selection event, leading to a negative phase shift relative to the selection itself.

The situation is quite different for the ancestral distribution. Because the fitness potential is the backward-time integral of the fitness, the mutational bias in the ancestors is strongest in the *preceding* season: mutations establishing the summer-adapted strain are most over represented in the spring, and for the winter-adapted strains in the fall. From the optimality perspective, the ancestral distribution shifts to summer-adaptation early to maximize the benefit of a summertime fitness advantage Figure 4.

## 4 Maximum likelihood inference on reconstructed trees

Suppose that we observe a genotype *G*_1_, *G*_2_ … in a populations at several sample times *T*_1_, *T*_2_ … and we can reconstruct the tree topology 𝒯 for the gene from linked neutral variation with minimal uncertainty. First we will calculate the exact likelihood and describe the path to its complete evaluation. Then we will show how the weak result allows the calculation of the likelihood in a more tractable form.

### 4.1 Exact likelihood particle methods

We construct the exact likelihood given a particular frequency history *X*, a path through frequency space governed by the Fisher-Kimura stochastic generator *L*.

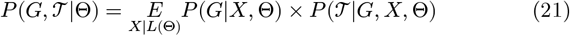

Here, 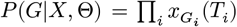 represents multinomial sampling at each time point. The difficult term is the expectation 𝔼_*X*|*L*(Θ)_, which requires integrating over a vast, high-dimensional space of possible frequency paths. Evaluating this integral is computationally intractable for realistic datasets, often requiring expensive particle filter methods (see subsection C.3 for a detailed formulation).

### 4.2 Weak-selection likelihood

In the weak-selection limit, we can leverage the frequency independence to dramatically reduce the effort required to evaluate the likelihood of genotypic data. Instead of using Monte Carlo sampling to estimate the integral over *X*, we can analytically marginalize out these hidden paths.

We begin by separating the probability of the tree topology and the probability of the data, given the tree.

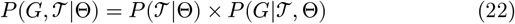

The tree likelihood is (in the same weak-limit) well approximated by the neutral coalescent, with lineage independent hazard 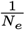. For coalescent times *t*_*c*_,

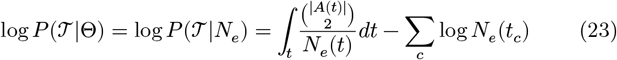

Where *A*(*t*) is the set of lineages alive at time *t*, and 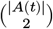 counts the number of interacting pairs.

To evaluate the data likelihood *P* (*G*|𝒯, Θ), we must account for the interactions between lineages due to coalescence hazard. Derived in subsection C.1, coalescence induces an interaction term in the fitness potential *ψ*^*i*^ of lineage *i*, This term combines with the single-lineage dynamics to yield lineage-coupled ODEs:

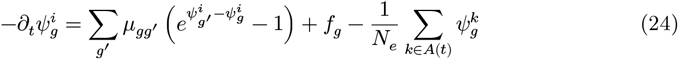

At the end of a lineage *T*_*i*_, where children *C*(*i*) coalescen into a parent *i*, we have the boundary condition

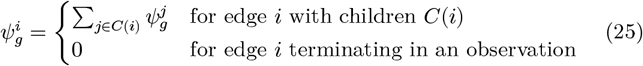

The data likelihood can be computed efficiently using a standard recursive pruning algorithm. The fitness potential defines a time-dependent, biased mutational operator 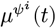 Equation 13 along each branch *i*. These 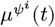 are the generator for the Kolmogorov equation, integrated backward from the tip:

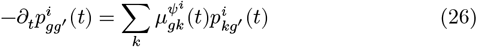

with the boundary condition at the tip, 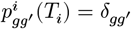. The likelihood is then found recursively from the tips to the root:

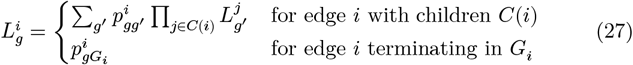

Then the total probability of th eobservations given the tree is

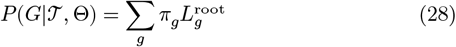

where π is a chosen prior (see subsection C.2) over the common-ancestor state.

### 4.3 Inference of seasonal parameters

We can infer the parameters of the seasonal selection model on trees generated from Monte Carlo population simulations. We find that we can collapse the phase parameter and infer the amplitude of the selection even for moderately-weak selection pressure Figure 5, even though the timeaveraged fitness difference between the alleles is zero and the mutation rate is low (*θ* = 0.1). The bias due to our weak-selection theory distorts some of the absolute values of the inference parameters from their true values in the simulation.

**Figure 5.**
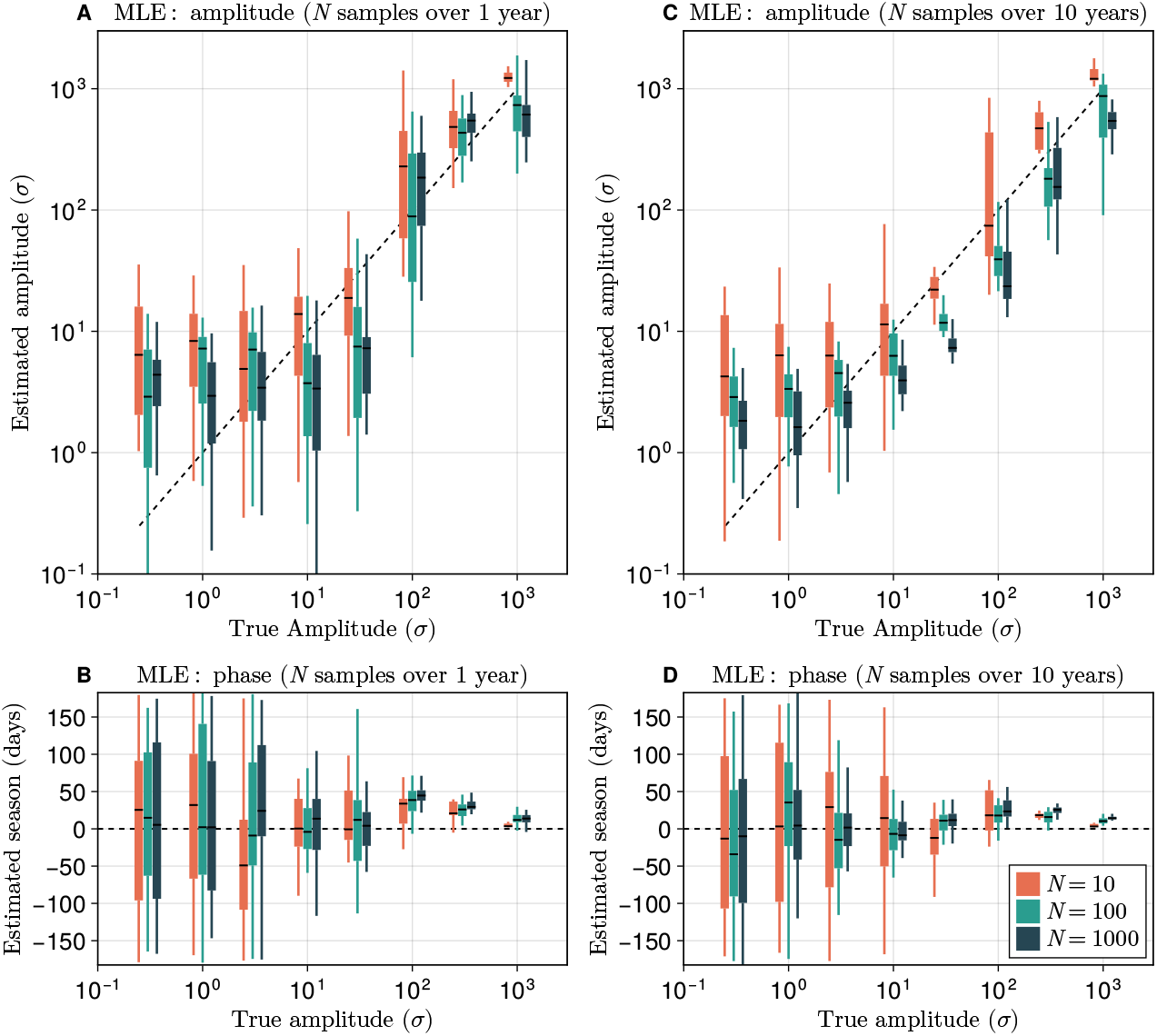
Inferring amplitude and phase of seasonal selection. We plot the distribution of maximum likelihood estimators (MLEs) for the amplitude and phase of the seasonal selection model where true parameters are *a* = 2/*N*_*e*_, and *φ* = 0. The population size is 10^3^, and the diversity parameter is *θ* = 0.1 for 10, 100 and 1000 samples. On the left (right) the observations are spread over a single year (10 years). Overfitting biases the inferred amplitude high (especially with only 10 samples N=10), and fit a random phase. At long obersrvation time and strong-selection the phase inference undergoes a transition and collapses near the true value. However, distortions from the weak-selection approximation generates a small phase bias and leads us to somewhat underinfer selection even with *N* = 1000.

Comparing the 1 year observational span and the 10 year span, increasing the sampling timeframe is comprable to increasing number of observations in terms of decresing the maximum-likelihood estimator (MLE) variance and resistance to overfitting. This is because spacing observations out further gives us more effectively independent seasons in which to see the one allele takeover the other.

## 5 Discussion

### Anticipatory dynamics

It seems paradoxical that a lineage, like an optimal actor, anticipates a future environment, but this is the same paradox that underlies the survivorship bias. The lineages we are concerned are conditioned on surviving to the observation time *T*.

It is instructive to consider a recent strong selection event in our own evolutionary past, the Black Death. Mutations that increased the expression of ERAP2 allowed more efficient presentation of bacterial peptides and conferred a survival advantage after the introduction of the plague in Europe in 1347 CE [28]. Tracing back the family tree of the survivors after this first wave of plague, the survivorship bias would create the impression of a coordinated aquisition of ERAP2-enhancing mutations [7] in an exponentially increasing bias up until (1347 CE). If we imagine viewing the complete record of meiotic recombination and point substitutions along the lineages of these survivors, it would appear as if these ancestors were *preparing* for the Black Death at the molecular level (see Figure 3).

This kind of “preparation” has little in common with an agent acting based on knowledge of the future. These ancestors of survivors could not foresee benefits of potential ERAP2 mutations. What we aim to highlight with these analogies is that the lineages and our dual control agents are statistically indistiguishable at the path level. Via this duality, reasoning about why an agent might take a certain path can provide intuition and predictive insight into why mutations are distributed as they are on successful lineages.

### 5.1 Relation to existing theories

### 5.1.1 Current approaches to phylogentic inference

Inference strategies generally fall into two categories: “tree methods” and “trajectory methods.” Despite their respective limitations, both approaches possess technical merits that make them appropriate for state-of-the-art inference. [45, 6, 33].

Tree methods typically employ a birth-death-sampling model [47, 30]. The central trick of these models ([35]) is that if we keep track of the probability of an individual *not* being sampled at any future time, the birth-death model *P* (*G*, 𝒯|Θ(*X*)) becomes directly integrable via Felsenstein pruning. However, to maintain this tractability, standard implementations typically treat lineages as conditionally independent entities governed by piecewise-constant rates (e.g., skyline models). This decoupling introduces two fundamental limitations. First is over-parameterization: by ignoring the mechanistic constraint that couples growth rates to the population state 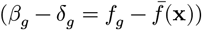, these models require an explosion of free parameters to describe dynamics that are physically generated by a small number of selection coefficients acting on a changing background. Second, and importantly for endemic populations, is the noise structure. By treating lineages as independent branching processes, these methods implicitly assume the variance structure of an infinite population (where variance scales with birth rate). They therefore fail to capture the information decay inherent to finite populations (where variance scales with drift, ~ 1/*N*_*e*_), potentially overestimating the confidence of inference in the deep past (see Appendix D).

Trajectory methods, in contrast, largely ignore the tree topology to study the frequency paths 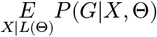 directly. Here, fitness effects are observed via their impact on frequency changes in accordance with the replicator dynamics, 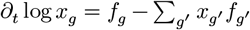 [24, 42, 46, 44, 43, 16, 34, 31]. While the full Kimura operator *L*_**x**_ itself induces an all-but-intractable PDE for its propagator, efficient approximations allow for inference at scale. However, these methods ignore the specific coalescent structure implied by the observed linked variation. Even when exact, trajectory methods “leave likelihood on the table”, extracting only the parameter information available in marginal allele frequencies while discarding the rich signal contained in the genealogy itself. This lost signal is not just lost precision; the tree structure carries the covariation information that resolves the difference between true selection and selection on a linked allele (genetic draft).

Our method bridges this divide: it utilizes the full resolving power of the tree structure (like birth-death models) but accounts for the lineage-population coupling and finite-population drift (like replicator dynamics) via the fitness potential. Via Equation 4, the correlation between the lineage state and the population state is explicitly handled through the backward Kolmogorov evolution *L*_**x**_, giving us the correct noise-structure.

An advantage of the fitness potential is its scalability to high-dimensional genotype spaces. In standard multi-type birth-death models, inference on sequence data requires tracking the joint distribution of all sites to calculate rates, a task that scales exponentially with sequence length (*G*^*L*^). Consequently, these methods must coarse-grain sequences into arbitrary “clades.” In contrast, our fitness potential factorizes under an additive fitness model. If we assume additive fitness (independent sites), the total potential decomposes into a sum of site potentials:

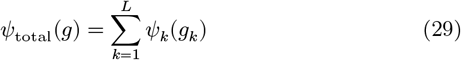

Consider a mutation event that occurs only at site *j*, changing genotype *g* → *g*′. Since *ψ*_*g*′_ and *ψ*_*g*_ are identical at all sites except *j*, the difference in potential—which drives the transition probability—depends only on the local contribution:

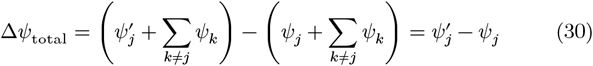

The probability of a transition at site *j* therefore depends only on the fitness potential at site *j*, with the complex background interactions absorbed into the shared normalization. This decomposition allows for sitespecific fitness inference at scale.

#### 5.1.2 The landscape of reproductive value

Our fitness potential Ψ(**x**, *t*) generalizes Fisher’s concept of reproductive value to dynamic, non-equilibrium populations [17]. In the limit of constant fitness and infinite time horizons, Ψ converges to the logarithm of the dominant left eigenvector of the evolutionary generator—exactly recovering Fisher’s classical definition. However, for time-varying selection (such as seasonal adaptation) or finite observation windows, Ψ captures the instantaneous reproductive value: a time-dependent field that guides lineages not toward a static optimum, but toward survival at the specific sampling time *T*.

The standard fitness landscape *f*(**x**) describes the instantaneous forces (birth and death rates) acting on the population. In contrast, the fitness potential Ψ(**x**) describes the integrated value of a state. In physics terms, if fitness *f* acts as a local force or gradient driving frequency changes, then Ψ acts as the potential energy surface. By transforming the problem from the local landscape (*f*) to the global potential landscape (Ψ), we move from describing the causes of evolution (selection) to describing the local maximization that takes place along the ancestral lineages.

#### 5.1.3 Relationship to other evolutionary path integrals and quasispecies theory

For the dual control problem with infinite time horizon (*γ* = 0), the exponential of the expected reward 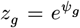 at time *t* can be solved by a linear ODE to the terminal time (the Bellman equation), *T* for which the cost function is zero afterwards. The linear ODE can be represented by a path integral.

The undiscounted HJB equation is related to the quasispecies operator (*μ* + *f*)_*gg*_′ by the Feynman-Kac lemma. This same path integral has been studied by previous authors [1, 9, 2]. Our focus is on different set of population statistics than the quasispecies theory. As a forward-in-time operator, the quasispecies equation acts on the vector of genotypic log-frequencies log *x*_*g*_ [9, 5, 12, 53]. In contrast, the dynamics of *ψ* runs backward in time, Equation 41, and is observable through genotypic *transitions* along a lineage (Equation 13) rather than in the frequencies of the population *x*_*g*_ itself, which we have decoupled entirely from the dynamic picture in the weak-selection limit. The quasispecies is an infinite population limit. The fitness potential carries the effects of the finite population via the discount rate *γ*^−1^ = *N*_*e*_. This discounting reflects the loss of *future* information backward-in-time.

We briefly note that the Wright-Kimura equation itself defines a different path integral at the population level that is closely related to the path integrals of non-equilibrium thermodynamics [36, 37]. The lineage path integral operates over the space of genotypes while the population path integral operates over *distributions* of genotypes.

### 5.2 Limitations

#### Dependence on tree data

The phylogenetic trees used as input are not directly observable but must be reconstructed from sequence data. While tree inference is computationally challenging, significant progress in maximum-likelihood and rapid heuristic methods has made accurate reconstruction feasible even for massive datasets [41, 52, 29, 15, 18, 39]. In this work, we condition on a fixed tree topology, an approach appropriate for the “big data” regime where the phylogenetic signal is strong and topological uncertainty is subordinate to demographic stochasticity.

However, the framework is not mathematically limited to fixed topologies. The fitness-potential likelihood *P*(𝒯|Θ) derived here constitutes a rigorous probability density for the tree structure under selection and finite-population drift. It is therefore suitable for integration into Bayesian phylogenetic samplers (e.g., BEAST2, RevBayes [32, 22, 6]), serving as a physics-aware drop-in replacement for standard coalescent or birth-death priors. This integration would allow future implementations to marginalize over topological uncertainty while retaining the correct mechanistic handling of lineage competition.

#### Weak selection and model robustness

The most biologically interesting regime, where selection is very strong, breaks the assumption at the heart of our derivation. Despite this, after suitable averaging over **x** with the equilibrium distribuiton, the frequency dependence often decouples so that Ψ_*g*_(*x*) = *ψ*_*g*_ − *φ*(*x*) also for arbitrarily strong selection. The obvious question remains: How strong is too strong?

While our derivation formally relies on the weak-selection limit, the method exhibits generality beyond this regime. Because the replicator dynamics are intrinsic to the formulation, the inference remains monotonic with respect to selection strength (see Results). In the strong selection limit, the primary deviation is an unknown rate of attenuation in the inferred signal. However, in practice, this attenuation is mathematically indistinguishable from deviations in the true effective population size, *N*_*e*_, which itself depends on the method of measurement and detailed population structure.

We contend that the approximation of the fitness potential is likely robust compared to the fundamental assumptions of the underlying Fisher-Kimura diffusion. The diffusion limit assumes all alleles are always present (ignoring stochastic extinction boundaries), and ignores spatial structure and complex frequency-dependent selection. These standard model approximations likely introduce more significant deviations from biological reality than the weak-selection decoupling at the heart of our method. Given the potential for model mismatch in real biological data, we emphasize that any inference must be statistically calibrated. The distribution of likelihoods obtained from the neutral component of the genome should serve as the null baseline to determine the statistical significance of inferred selection parameters.

## 6 Conclusions

A remarkable link between control theory and evolution has been previously proposed [49, 26, 50]. This link has been used to motivate advanced control theory sampling methods and is related to state-of-the-art deep learning methods such as stochastic tree-search. We have shown that this link can also be reversed to derive a quantitative statistical theory for the ancestral proccess. Based on this theory, the fitness inference method we present can be applied to time dependent fitness models, out of equilibrium. We can infer both positive and negative selection, and we do not require inferring an underlying genotype frequency **x**. The method makes direct use of linked neutral variation, which is represented by the temporal embedding of the reconstructed tree. However, detailed frequency dependence reemerges in the strong-selection clonally interfering limit, making the method only approximate for important class of problems: strong selection on large effective population sizes over significant timescales.

Selection on inherited traits does not specifically require genetic material, and inherited epigenetic states transmitted between generations, or correlated environmental conditions between parent and child could likewise be treated with a control theoretic framework. Evolution and control are linked at a level more fundamental than genetic inheritance itself.

The lens of optimal control theory provides a fascinating perspective on the apparent emergence of adaptive planning. The apparent “stratgies” emerging from Darwinian adaptation can now be mathematically mapped precisely to optimal strategies in a corresponding game of stochastic control. An evolving pathogen appears to be an opponent playing optimally against human mitigation strategies because this optimal play (within the problem constraints) is a general emergent feature of Darwinian adaptation from the perspective of the common ancestral line.

## 7 Declarations

### 7.1 AI Usage Statement

Large Language Models (Gemini 2.5, Gemini 3, and ChatGPT 4.1o) were used during the drafting process to assist with iterative editing, refining the narrative, and checking for clarity for a broad audience. The mathematical derivations, conceptual framework, and code implementation (except where noted) were generated by the author. All AI-suggested text and code was critically reviewed, verified, and revised by the author, who assumes full responsibility for the accuracy and integrity of the work.

## 7.2 Acknowledgements

The author would like to acknowledge important conversations with William de Witt, Jakub Otwinowski, and Armita Nourmohammad. Initial work was partially supported by CRC 1310 Young Researchers Grant.

## 7.3 Code Availability

Code for the fitness potential framework and figure generation is available at: https://github.com/chelate/fitness_potential_paper

## A The control theoretic view of evolving populations

To introduce the mathematical duality at the heart of our method consider the code for a population-genetic simulation, `popgen.c.` The program takes as inputs *N* and *λ*, and *f* and *μ*, as time varying functions. It then generates a population of size *N* and evolves it forward in time with birth and death rates

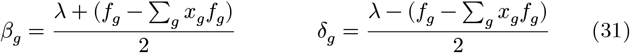

and mutation *μ*_*gg*_′ from time 0 to time *T* evaluated with the Gillespie method, keeping track of the lineage relations of the entire population. The same code can be viewed in two ways:

- A population geneticist sees a Moran-type Wright-Fisher simulation
- A control theorist sees a Monte Carlo algorithm for solving a continuous-time stochastic control problem.

In the following sections we reverse engineer the parameters of the control problem that `popgen` seems to be solving from the control-theorist perspective.

### A.1 Reverse engineering the control problem

We recall the basic principles of optimal control theory. One seeks policies *u*_*h*_(*g, t*), which determine our action *h* as a function of state *g* and time, so as to maximize expected rewards. Because of the general Markov property, optimal policy is a local function of the expected optimal reward at each state—the value function *ψ*(*g, t*). Therefore, constructing the value function amounts to solving the control problem.

#### Path-integral control

The value function satisfies the Bellman equation, a partial differential equation in continuous space. For (typical) highdimensional systems the Bellman equation is intractable. However, in the remarkable case of path-integral control, we can recast the non-linear Bellman equation into a linear partial differential equation [26, 49, 50]. This linear partial differential equation may then be estimated using a forward-time evolutionary process [11, 10, 25]. The evolutionary process targets the corresponding path integral in the Feynman-Kac lemma for the linear-transformed Bellman equation.

In this evolutionary process, the log-number of descendants in a particular state *g*, is then, in expectation, the optimal total reward (the value function) of the control problem. Since the value function is sufficient to reconstruct the optimal control, the solution to the control problem can be extracted from the expectations of this evolutionary process [48]. This biomimetic strategy for solving partial differential equations also appears in other fields e.g. particle filters for transfer-matrix methods in solid state physics [23].

### A.2 Entropy-regularized control on continuous time and discrete space

A control theorists therefore sees `popgen` as exactly this kind of stochastic solver. To be precise, `popgen` must be targeting the path integrals for a continuous time Markov jump process defined on genotypic space *G* with transitions governed by *μ*_*ij*_ i.e. a mutation process. We can infer that the controller in the target problem is able to exponentially-tilt the mutation rate into state *g*′ by a factor *u*_*g*_′ (*g*), so that the controlled rate of transitioning from *g* → *g*′ is

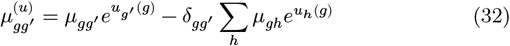

The reward rate of the dual problem is equal to the fitness function of the genotypic state *f*_*g*_(*t*), minus a KL-Divergence rate which characterizes the cost of controlling the system

We derive the Bellman equation for the dual control problem. This extends the discrete time case from [50] to continuous time Markov control. ^1^. We apply the discrete space and time results of [50] with transition function *p*(*g*′|*g*) = δ_*gg*_′ + *τ μ*_*gg*_′ and take the formal limit *τ* → 0.

#### Infinitessimal KL-divergence

Our continuous-time derivation of the appropriate cost deviates from [50] in that instead of enforcing normalization with a lagrange multiplier, we start from a normalized description of continuous time KL-control, where the rows of the generator sum to zero, implying that the conditional probability generated sums to one. The diagonal terms *u*_*gg*_ in the control cancel and are completely arbitrary:

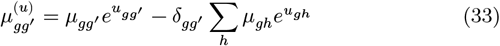

The KL divergence rate can be calculated by the short-time Taylor expansion of the matrix exponential

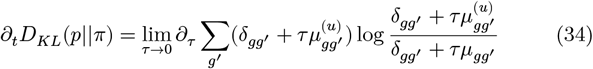

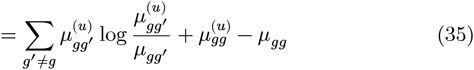

since *μ*_*gg*_ = − Σ_*h*≠*g*_ *μ*_*gh*_

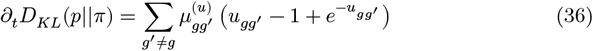

Thus the reward rate is

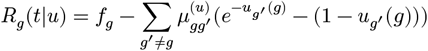

Typically *R* (and the terminal condition that control ends at *t* = *T*) can be seen as the fundamental specification of the control problem. Instead, we have reconstructed this function by working backwards from the sampling algorithm.

With the reward function *R* in hand, its straightforward to optimize the control and construct the HJB equation. The change in the total optimal expected reward (backward in time)

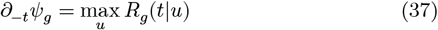

We determine the optimal control as a function of *ψ*

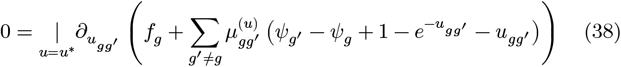

this simplifies to

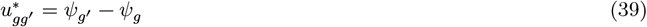

with the HJB equation becoming

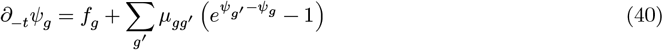

The optimally controlled mutation operator is therefore 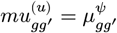, where,

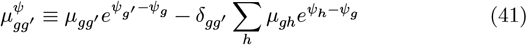

Comparing Equation 41 and Equation 13 jutifies the statement that ancestral lineages appear optimally controlled. This recapitualtes a remarkable result from [26] that surviving paths generated by following the parenthood relationships and view them as paths forward in time appear to be sampled from the ensemble of optimally controlled trajectories.

#### Comparison with the Pop-gen solution: Missing discount term

When we compare Equation 41 to Equation 12, we see an important difference: the result above Equation 41 does not include a discounting term. Inserting a discounting term 1/*N*_*e*_*ψ*_*g*_ where *N*_*e*_ = *N*/*λ* is a familiar modification of the HJB equation in control theory. A non-zero discount is essential for stability in reinforcement learning, with the discounting usually considered a necessary hyperparameter

However, this was *not* predicted by the path-integral control theory (PIC). Indeed, a discounting term breaks the assumptions upon which the path integral form depended, spoiling the linearity which allowed the use of the Feynman-Kac lemma. Our analysis from population genetics explain the emergence of a discount and in a sense can be seen as a finite-population modification of the Feynman-Kac lemma.

### A.3 HJB path integral and connection to forward evolutionary process

We can recast this equation as a linear equation via an exponential transformation.

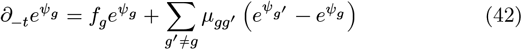

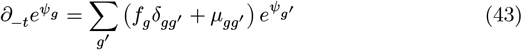

defining 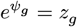 we recognize the linear quasispecies equation backward in time.

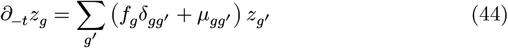

This can be represented by the path integral

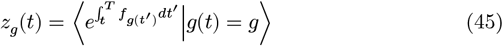

with path density determined by the neutral mutation generator *μ* and starting from *g*. By construction, *z* can be estimated up to a multiplicative constant via an evolutionary algorithm, where in the infinite population size limit, the average number of descendants of individuals in a genotypic class *g*, alive at time *T* is proportional to *z*_*g*_(*t*).

#### The path integral

Inserting a discrete time mutational operator at ever-finer intervals, we see that the number of descendants can be expressed as a path integral generated by the continuous mutational process,

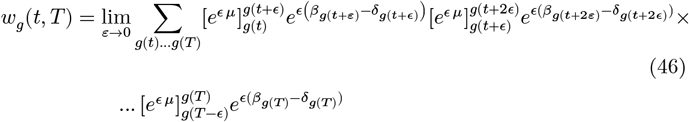

Where 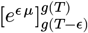 is the matrix exponential, (the short time propagator for the neutral mutation process).

#### Effect of discounting on path-integral control

We find that we can account for genetic drift of finite population sizes by adding a discount rate *γ* to the control problem. In [51] discounting is treated as a modification of the (in that reference, discrete-time) equation for *z*.

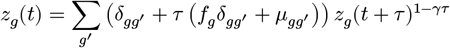

In the continuous limit this yields the modification,

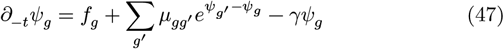

where we have determined that the discounting time scale and the coalescent time scale should match *γ* = 1/*N*_*e*_. Thus the optimal control remains the same, only the Bellman equation becomes modified by a decay factor.

## B Details of coordinate transform effect on the Kimura operator

The mutation operator 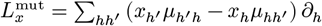

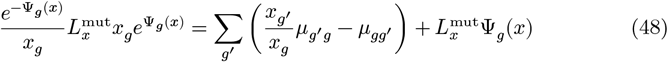

and 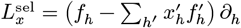

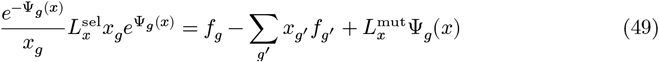

and finally 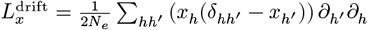

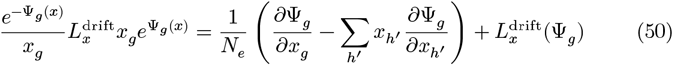

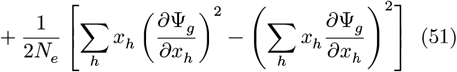

The end result of the coordinate transformation is

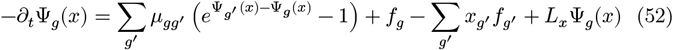

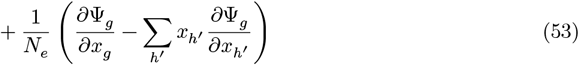

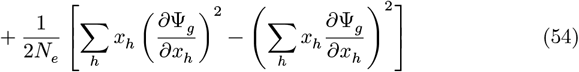

## C Inference-methods and tree likelohood

### C.1 Interactions between fitness potential of contemporaneous lineages

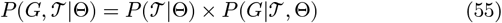

Our parameters are those neccesary for specifying the Kimura operator the (potentially time-dependent) drift, mutation, fitness functions Θ = *N*_*e*_, *μ, f*.

To apply our weak-selection results for this problem, we must determine how coalescence affects the fitness potential *ψ*. For two lineages *i* and *j* to coalesce in a short time *τ*, one of the *Nx*_*g*_*β*_*g*_*τ* coalescent event must occur on exactly this pair from the 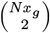 possible *g*-lineages in the population.

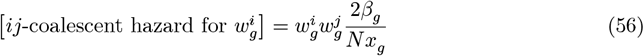

since we are concerned with the weak selection limit anyway, we can replace 2*β*_*g*_ /*N* = 1/*N*_*e*_, ignoring terms of size 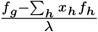. Using the same weak selection and gauge choice as in the single lineage case

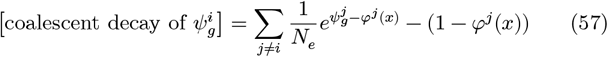

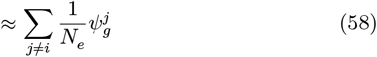

Including this interaction term into Equation 12 results in an equation for the evolution of *ψ*^*i*^ along each of the branches. These coalescent interactions now enforcee an enhanced decay along each of the branches *i* that coexist with other branches *k* at time *t*.

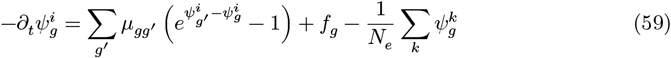

At the moment of coalescence *C*(*i*) → *i*, we have the boundary condition at *T*_*i*_

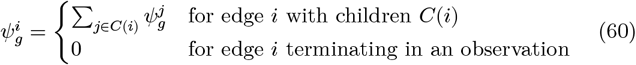

This decay term can be understood as the observed lack of coalescence drives decoherence in the fitness potentials of contemporaneous lineages.

#### Tree likelihood is neutral at same order

The tree likelihood is determined by the coalescent hazard Λ_*ij*_(*t*) for contemporaneous lineages *i j*, exactly as a Poisson process with time varying rates, with rate

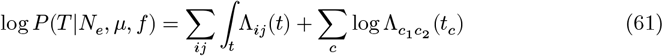

where the second sum is over the observed coalescent events *c* between lineages *c*_1_ and *c*_2_ occuring at time *t*_*c*_. To account for the fact that lineages must share a type to coalesce, making the hazard

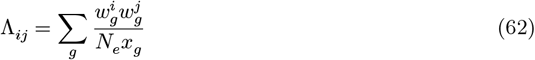

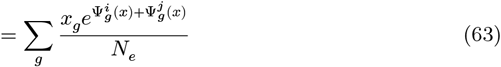

assuming the self-consistent decomposition as before 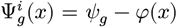, we have, to second order

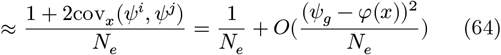

In the weak selection limit, the frequency dependence of the tree likelihood vanishes, and luckily the coalescent hazard appears neutral at the same approximation order as the weak selection result, 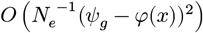.

In this case the tree likelihood is only a function of the effective population size *N*_*e*_. For coalescent times *t*_*c*_,

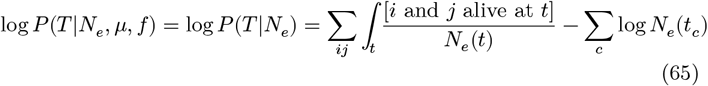

with [X] the Iverson bracket, 1, if X is true and 0 i X false. The cancelation in the denominator *x*_*g*_, reflects the fact that, under weak selection, tree structure is unaffected by allelic diversity and mutations can be trivially painted on a neutrally coalescent tree.

### C.2 Observation likelihood from the ancestral-biased mutational operator

Given functions *f*_*g*_ and *μ*_*gg*_′ and *N*_*e*_, these define via Equation 60 a time dependent fitness potential over the genotypic state space at each point along along each edge *i* of this tree Equation 60 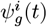, and a biased mutational operator *μ*^*ψ*^(*t*) for times between the lineage coalescence *t*_*i*_ and its termination in an observation or birth event at *T*_*i*_. The biased mutational operator *μ*^*ψ*^ then determines a conditional probability density for an initial genotype *g* to evolve into a new genotype *g*′ by the end of the lineage exactly analogous to Felsenstein’s pruning algorithm [14, 6],

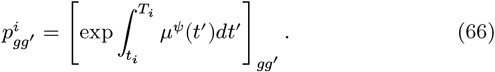

where exp *∫* ·*dt* is the time-ordered matrix exponential. The likelihood of observing a tree rooted on lineage *i* starting in state *g* is defined recursively

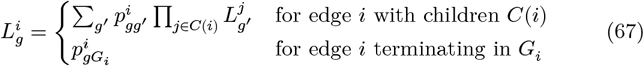

#### Common ancestor prior

We must choose an averaging scheme over the final common ancestor state. We propose the stationary distribution of the ancestral process at the root time as a natural prior 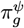 of 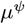 which is easy to compute.

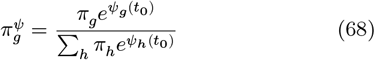

Then the total probability of the observations given the tree is

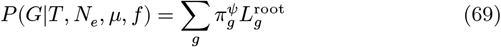

Other choices, such as a uniform prior, are also be possible.

### C.3 Exact inference using the lineage frequencies and marginalization over paths

For contrast, we briefly describe the procedure to construct the exact likelihood given a particular frequency history *X*, a path through frequency space governed by the Fisher-Kimura stochastic generator *L*^†^.

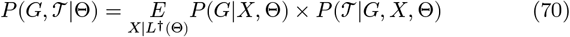

Here, 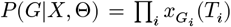 represents multinomial sampling at each time point.

With each proposal path we can evaluate the tree likelihood *P* (𝒯|*G, X*, Θ) using the *forward* (but still time reversed, still from tip to root!) Kolmogorov equation with *μ*^rev^(**x**) for each lineage **w** with coalescnce hazard described above

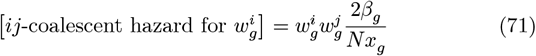

With sophisticated importance samplers for *X*, this exact likelihood might become feasible for small systems and few time points.

## D Derivation of a “birth-death potential” and noise mismatch

Recent state-dependent birth-death models [33] explicitly modify forward transition rates by the ratio of sampling probabilities 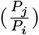, an operation mathematically equivalent to the optimal control policy derived in our framework (*μe*^ΔΨ^). A tree likelihood could theoretically be computed using the raw backward Kolmogorov equation, but these methods take an extra step of conditioning the forward generator on eventual sampling, finding that this conditioned process yields a more stable likelihood for inference. Our work provides the theoretical justification for this practice of using the forward equation: the raw likelihood is inappropriate for inference because it comes from the wrong (unconstrained, unconditioned) model. But the forward propagator is equivalent to a “birth-death potential” that steers lineages away from extinction, and is quite similar to our fitness potential.

However, this birth-death potential has distinct dynamics. We consider a standard multi-type birth-death process conditioned on a single sampling event. Let *E*_*g*_(*t*) be the probability that a single lineage currently in state *g* at time *t* will not be sampled (i.e., goes extinct or remains unsampled) before the present time *T*.

In standard birth-death-sampling models, this probability evolves according to the backward Kolomolgorov birth-death equation:

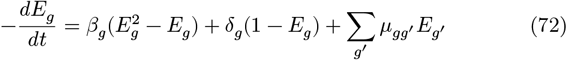

where *β*_*g*_, δ_*g*_, and *μ*_*gg*_′ are the birth, death, and mutation generator respectively. We define the survival probability, *P*_*g*_(*t*) = 1 − *E*_*g*_(*t*) that the lineage is sampled. This closure is only possible in the case of a single sample event, otherwise there are two separate equations. Substituting *E*_*g*_ = 1 − *P*_*g*_ into the Riccati equation and simplifying using the relations for net fitness 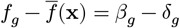,

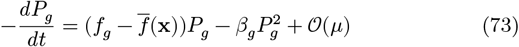

We apply the logarithmic transformation to map this probability to a fitness potential 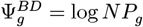. The boundary condition at sampling time *T* for a single identified lineage is *P*_*g*_(*T*) = 1/*N*, implying that 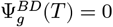 matching the fitness potential boundary condition.

We can therefore rewrite the non-linear decay term

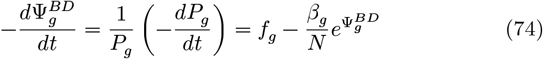

The effective population size in terms of the noise intensity is *N*_*e*_ = *N*/*λ*. Assuming weak selection (*f* ≪ *λ*), the birth rate is dominated by turnover, *β*_*g*_ ≈ *λ*/2. We can therefore rewrite the non-linear decay term:

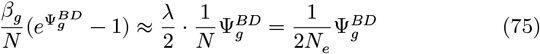

Substituting this back into the evolution equation:

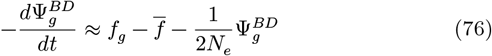

Comparing this result to the fitness potential derived from the WrightFisher process Equation 12, which contains a decay term of 1/*N*_*e*_, we see a discrepancy of a factor of 1/2 in the decay term despite them targeting the same continuum process. This birth-death potential decay term is in contradiction with numerical experiments and the analytic results for the rare-mutation steady-state limit of a biallelic system (see subsection 3.1).

## E Additional models

### E.1 Triallelic system

For completeness we consider more complicated systems beyond the biallelic case. Since mutations are relatively rare, the fitness potential exponentially decays backward in time. We sample the population at time at 2*N*_*e*_ and then imagine we can reconstruct the population tree with perfect fidelity. We trace a lineage backward in time and record the genotypic changes. We compare these observations to the predictions from the optimal control result on the dual control problem as shown in Figure 6. In this simulation we consider three genotypes of differing fitnesses.

**Figure 6.**
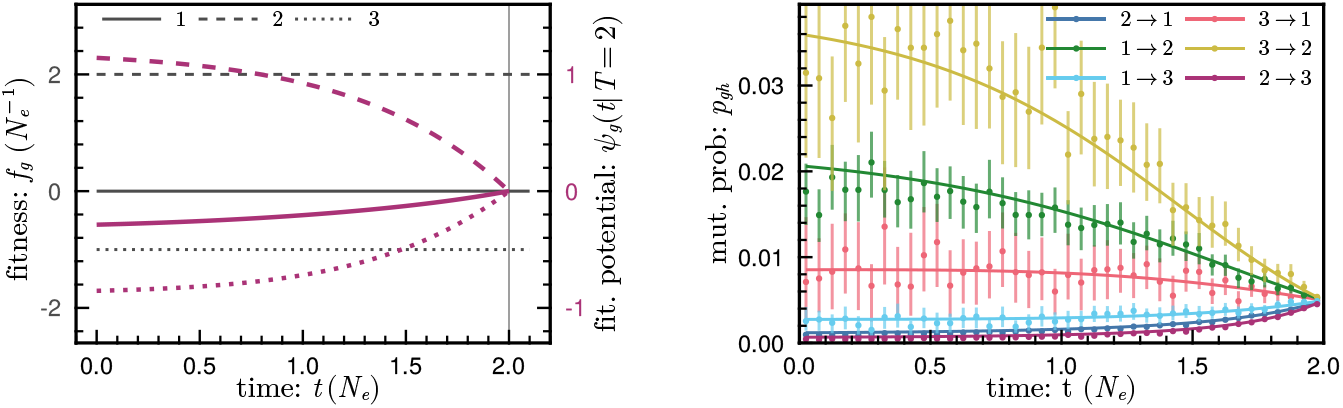
Constant fitness. The lineage path is the path traces through genotypic space going backward in time along a lineage, it appears to obey a biased mutational operator *μ*^*ψ*^ governed by *ψ*. Near the sample time 2*N*_*e*_, the mutations approach their neutral values, but looking backward in time they approach their equilibrium values.

**Figure 7.**
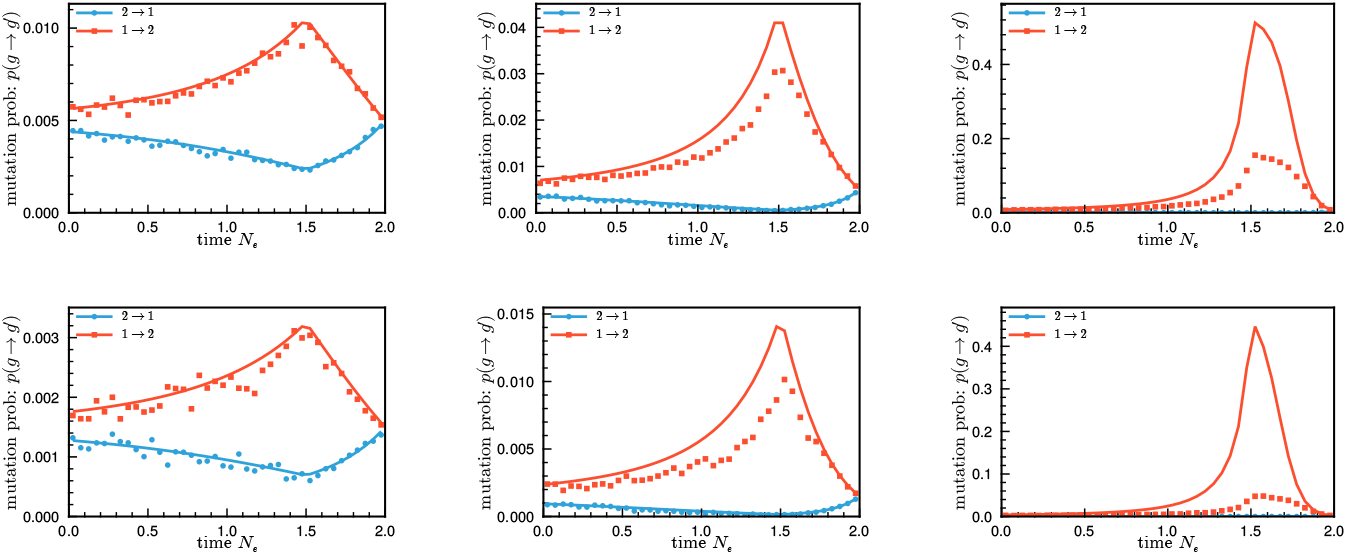
Breakdown of theory with strong positive selection. Shown are the results for the biallelic pulsed fitness model (between 1.5 and 2.0 *N*_*e*_) sampled at 2.0*N*_*e*_. When mutation rates are low (upper panels *θ* = 0.01lower panels *θ* = 0.03) and selection is stronger (left to right), we see that theory predicts more mutational biasing than actually occurs. The emergence of a new positively selected mutant is strongly correlated with an increase in the mean fitness of the population, and this leads to stiffer competition due to the self interatctions.

The fitness potential is the time integral of these fitness differences backward in time.

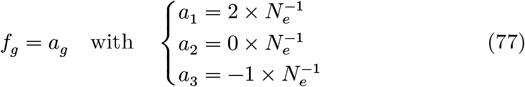

### E.2 Breakdown of theory with strong positive selection and low mutation rates

At the mutational level there is a breakdown in the symmetry of the mutation rate bias around the neutral value. When plotted on a log scale we can see that even for the case where the breakdown is worst, there is a significant amount of bias in the amount of positive selection.

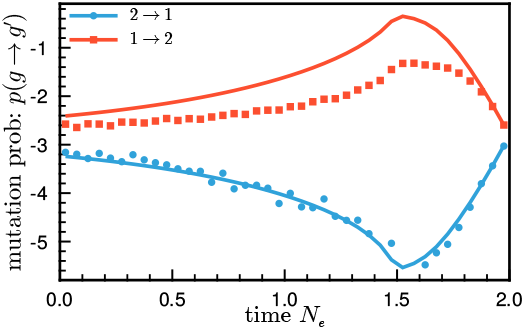

Suppose that a mutation is positively selected to the extent that every time there is a mutation in the population, it reaches fixation. The rate of such events occurring is only *μN*, clearly the maximum control on a positively selected allele is Δ*ψ*_max_ = log *N*. On the other hand, for negative selection, there is no limit to how often we can reject a given mutation. This kind of breakdown can therefore be seen as a result of the discrete nature of the population relative to the continuum limit of the Kimura equation itself.

## F Simulation details

Simulations were performed in Julia [4] using a balanced Moran process with units *N* = 1000, and *λ* = 1 giving *N*_*e*_ = 1000 in some arbitrary units of time (in the seasonal model this is in days). The inference problem was simulated *N* = 2000 in the hope that this would lead to less biased results than *N* = 1000 (it didn’t). Unless otherwise specified the mutations are set so that the unitless parameter, *θ* = 0.1. Lower *θ* leads to larger deviations from the constant *N*_*e*_ theory due strong correlation in the fates of individual mutations and the average fitness of the population. The fitness potential differential equations were solved using the DifferentialEquations.jl package [40], MLE optimization was performed with the BOBYQA algorithm, [38] using the package PRIMA.jl, [54], plots made with Makie.jl [8].

## Footnotes

1 The entropic part of the reward function is the cost of control. This quantity can be interpreted as the rate at which we remove paths from an ensemble of paths obeying *μ* so that the sub-ensemble obeys *μ*^(*u*)^. In entropy-regularized or soft control, the entropy typically has a temperature parameter [21] which rescales the entropy cost. However, by using units of Malthusian fitness, we have implicitly set the temperature parameter to one.

## References

[1] Mark Ancliff and Jeong-Man Park. Spin coherent state representation of the crow-kimura and eigen models of quasispecies theory. J. Stat. Phys., 143(4):636–656, May 2011.

[2] Ellen Baake and Frederic Alberti. Solving the selection-recombination equation: Ancestral lines under selection and recombination. arXiv [math.PR], March 2020.

[3] Nicholas H Barton and Alison M Etheridge. The relation between reproductive value and genetic contribution. Genetics, 188(4):953– 973, August 2011.

[4] Jeff Bezanson, Alan Edelman, Stefan Karpinski, and Viral B Shah. Julia: A fresh approach to numerical computing. SIAM Rev., 59(1):65–98, January 2017.

[5] CK Biebricher and M Eigen. What is a quasispecies? Curr. Top. Microbiol. Immunol., 299:1–31, 2006.

[6] Remco Bouckaert, Joseph Heled, Denise Kühnert, Tim Vaughan, Chieh-Hsi Wu, Dong Xie, Marc A Suchard, Andrew Rambaut, and Alexei J Drummond. BEAST 2: a software platform for bayesian evolutionary analysis. PLoS Comput. Biol., 10(4):e1003537, April 2014.

[7] Graham E Budd and Richard P Mann. History is written by the victors: The effect of the push of the past on the fossil record. Evolution, 72(11):2276–2291, November 2018.

[8] Simon Danisch and Julius Krumbiegel. Makie.jl: Flexible highperformance data visualization for julia. J. Open Source Softw., 6(65):3349, September 2021.

[9] Esteban Domingo, Julie Sheldon, and Celia Perales. Viral quasispecies evolution. Microbiol. Mol. Biol. Rev., 76(2):159–216, 2012.

[10] A Doucet, N De Freitas, and NJ Gordon. Sequential monte carlo methods in practice. Springer.

[11] A Doucet, N Freitas, and N Gordon. An introduction to sequential monte carlo methods. Sequential Monte Carlo methods in, 2001.

[12] Manfred Eigen, John McCaskill, and Peter Schuster. Molecular quasi-species. J. Phys. Chem., 92(24):6881–6891, December 1988.

[13] Paul Fearnhead. The common ancestor at a nonneutral locus. J. Appl. Probab., 39(1):38–54, March 2002.

[14] J Felsenstein. Evolutionary trees from DNA sequences: a maximum likelihood approach. J. Mol. Evol., 17(6):368–376, 1981.

[15] Xizhou Feng, Kirk W Cameron, and Duncan A Buell. PBPI: a high performance implementation of bayesian phylogenetic inference. In SC ’06: Proceedings of the 2006 ACM/IEEE Conference on Supercomputing, pages 40–40. ieeexplore.ieee.org, November 2006.

[16] Andrew L Ferguson, Jaclyn K Mann, Saleha Omarjee, Thumbi Ndung’u, Bruce D Walker, and Arup K Chakraborty. Translating HIV sequences into quantitative fitness landscapes predicts viral vulnerabilities for rational immunogen design. Immunity, 38(3):606–617, 2013.

[17] R Fisher and JH Bennett. The genetical theory of natural selection: a complete variorum edition. 1999.

[18] Tomáš Flouri, Xiyun Jiao, Bruce Rannala, and Ziheng Yang. Species tree inference with BPP using genomic sequences and the multispecies coalescent. Mol. Biol. Evol., 35(10):2585–2593, October 2018.

[19] Daniel Goodman. Optimal life histories, optimal notation, and the value of reproductive value. Am. Nat., 119(6):803–823, June 1982.

[20] Alan Grafen. A theory of fisher’s reproductive value. J. Math. Biol., 53(1):15–60, July 2006.

[21] Tuomas Haarnoja, Aurick Zhou, Kristian Hartikainen, George Tucker, Sehoon Ha, Jie Tan, Vikash Kumar, Henry Zhu, Abhishek Gupta, Pieter Abbeel, and Sergey Levine. Soft actor-critic algorithms and applications. arXiv [cs.LG], December 2018.

[22] Sebastian Höhna, William A Freyman, Zachary Nolen, John P Huelsenbeck, Michael R May, and Brian R Moore. A bayesian approach for estimating branch-specific speciation and extinction rates. bioRxiv, February 2019.

[23] Yukito Iba. Population monte carlo algorithms. Trans. Jpn. Soc. Artif. Intell., 16(2):279–286, 2001.

[24] Christopher J R Illingworth, Jayna Raghwani, David Serwadda, Nelson K Sewankambo, Merlin L Robb, Michael A Eller, Andrew R Redd, Thomas C Quinn, and Katrina A Lythgoe. A de novo approach to inferring within-host fitness effects during untreated HIV-1 infection. PLoS Pathog., 16(6):e1008171, June 2020.

[25] Brian Jefferies. Evolution Processes and the Feynman-Kac Formula. Springer Science & Business Media, March 2013.

[26] HJ Kappen. Path integrals and symmetry breaking for optimal control theory. J. Stat. Mech., 2005(11):P11011, November 2005.

[27] M Kimura. On the probability of fixation of mutant genes in a population. Genetics, 47(6):713–719, June 1962.

[28] Jennifer Klunk, Tauras P Vilgalys, Christian E Demeure, Xiaoheng Cheng, Mari Shiratori, Julien Madej, Rémi Beau, Derek Elli, Maria I Patino, Rebecca Redfern, Sharon N DeWitte, Julia A Gamble, Jesper L Boldsen, Ann Carmichael, Nükhet Varlik, Katherine Eaton, Jean-Christophe Grenier, G Brian Golding, Alison Devault, Jean-Marie Rouillard, Vania Yotova, Renata Sindeaux, Chun Jimmie Ye, Matin Bikaran, Anne Dumaine, Jessica F Brinkworth, Dominique Missiakas, Guy A Rouleau, Matthias Steinrücken, Javier Pizarro-Cerdá, Hendrik N Poinar, and Luis B Barreiro. Evolution of immune genes is associated with the black death. Nature, 611(7935):312–319, November 2022.

[29] Sudhir Kumar, Koichiro Tamura, and Masatoshi Nei. MEGA3: Integrated software for molecular evolutionary genetics analysis and sequence alignment. Brief. Bioinform., 5(2):150–163, June 2004.

[30] Denise Kühnert, Tanja Stadler, Timothy G Vaughan, and Alexei J Drummond. Phylodynamics with migration: A computational framework to quantify population structure from genomic data. Mol. Biol. Evol., 33(8):2102–2116, August 2016.

[31] Colin LaMont, Jakub Otwinowski, Kanika Vanshylla, Henning Gruell, Florian Klein, and Armita Nourmohammad. Design of an optimal combination therapy with broadly neutralizing antibodies to suppress HIV-1. Elife, 11, July 2022.

[32] Giovanni Laudanno, Bart Haegeman, Daniel L Rabosky, and Rampal S Etienne. Detecting lineage-specific shifts in diversification: A proper likelihood approach. Syst. Biol., 70(2):389–407, February 2021.

[33] Maya A Lewinsohn, Trevor Bedford, Nicola F Müller, and Alison F Feder. State-dependent evolutionary models reveal modes of solid tumour growth. Nat Ecol Evol, March 2023.

[34] Raymond H Y Louie, Kevin J Kaczorowski, John P Barton, Arup K Chakraborty, and Matthew R. McKay. Fitness landscape of the human immunodeficiency virus envelope protein that is targeted by antibodies. Proc. Natl. Acad. Sci. U. S. A., 115(4), January 2018.

[35] Wayne P Maddison, Peter E Midford, and Sarah POtto. Estimating a binary character’s effect on speciation and extinction. Syst. Biol., 56(5):701–710, October 2007.

[36] Ville Mustonen and Michael Lässig. Molecular evolution under fitness fluctuations. Phys. Rev. Lett., 100(10):108101, March 2008.

[37] Ville Mustonen and Michael Lässig. Fitness flux and ubiquity of adaptive evolution. Proc. Natl. Acad. Sci. U. S. A., 107(9):4248– 4253, March 2010.

[38] MJ Powell. The BOBYQA algorithm for bound constrained optimization without derivatives. Cambridge NA Report NA2009/06, University of Cambridge, Cambridge, 26:26–46, 2009.

[39] Morgan N Price, Paramvir S Dehal, and Adam P Arkin. FastTree 2– approximately maximum-likelihood trees for large alignments. PLoS One, 5(3):e9490, March 2010.

[40] Christopher Rackauckas and Qing Nie. DifferentialEquations.jl – a performant and feature-rich ecosystem for solving differential equations in julia. J. Open Res. Softw., 5(1):15, May 2017.

[41] Pavel Sagulenko, Vadim Puller, and Richard A Neher. Tree-Time: Maximum-likelihood phylodynamic analysis. Virus Evol, 4(1):vex042, January 2018.

[42] David Seifert, Francesca Di Giallonardo, Karin J Metzner, Huldrych F Günthard, and Niko Beerenwinkel. A framework for inferring fitness landscapes of patient-derived viruses using quasispecies theory. Genetics, 199(1):191–203, 2015.

[43] Karthik Shekhar, Claire F Ruberman, Andrew L Ferguson, John P Barton, Mehran Kardar, and Arup K Chakraborty. Spin models inferred from patient-derived viral sequence data faithfully describe HIV fitness landscapes. Phys. Rev. E Stat. Nonlin. Soft Matter Phys., 88(6):062705, December 2013.

[44] Kai S Shimagaki and John P Barton. Efficient epistasis inference via higher-order covariance matrix factorization. bioRxiv, page iyaf118, October 2024.

[45] Kai S Shimagaki, Rebecca M Lynch, and John P Barton. Parallel HIV-1 fitness landscapes shape viral dynamics in humans and macaques that develop broadly neutralizing antibodies. Elife, 14(RP105466):RP105466, November 2025.

[46] Muhammad Saqib Sohail, Raymond H Y Louie, Zhenchen Hong, John P Barton, and Matthew R McKay. Inferring epistasis from genetic time-series data. Molecular Biology and Evolution, 39(10), October 2022.

[47] Tanja Stadler, Denise Kühnert, Sebastian Bonhoeffer, and Alexei J Drummond. Birth-death skyline plot reveals temporal changes of epidemic spread in HIV and hepatitis C virus (HCV). Proc. Natl. Acad. Sci. U. S. A., 110(1):228–233, January 2013.

[48] Evangelos Theodorou, Jonas Buchli, and Stefan Schaal. Learning policy improvements with path integrals. In Yee Whye Teh and Mike Titterington, editors, Proceedings of the Thirteenth International Conference on Artificial Intelligence and Statistics, volume 9 of Proceedings of Machine Learning Research, pages 828–835, Chia Laguna Resort, Sardinia, Italy, 2010. PMLR.

[49] Evangelos A Theodorou. Iterative path integral stochastic optimal control: Theory and applications to motor control. University of Southern California, 2011.

[50] Emanuel Todorov. Linearly-solvable markov decision problems. Adv. Neural Inf. Process. Syst., 19, 2006.

[51] Emanuel Todorov. Efficient computation of optimal actions. Proc. Natl. Acad. Sci. U. S. A., 106(28):11478–11483, July 2009.

[52] Yves Van de Peer and Marco Salemi. Phylogenetic inference based on distance methods. The phylogenetic handbook, pages 142–160, 2009.

[53] Claus O Wilke. Quasispecies theory in the context of population genetics. BMC Evol. Biol., 5:44, August 2005.

[54] Z Zhang. PRIMA: reference implementation for powell’s methods with modernization and amelioration. 2023.

